# A striosomal accumbens pathway drives compulsive seeking behaviors through an aversive Esr1+ hypothalamic-habenula circuit

**DOI:** 10.1101/2024.10.11.617042

**Authors:** Thomas Contesse, Buse Yel Bektash, Marta Graziano, Chiara Forastieri, Alessandro Contestabile, Salome Hahne, Felix Jung, Ifigeneia Nikolakopoulou, Xiao Cao, Vasiliki Skara, Ioannis Mantas, Sarantis Giatrellis, Marie Carlén, Rickard Sandberg, Daniela Calvigioni, Konstantinos Meletis

## Abstract

The lateral hypothalamic area (LHA) integrates external stimuli with internal states to drive the choice between competing innate or value-driven motivated behaviors. Projections from the LHA to the lateral habenula (LHb) shape internal states, with excitatory estrogen receptor 1-expressing (Esr1+) LHA-LHb neurons driving aversive responses and sustained negative states. Here, we identify and functionally characterize a specific projection from the nucleus accumbens (ACB) that targets Esr1+ LHA-LHb neurons. Using cell-type-specific tracing of monosynaptic inputs, single-nucleus RNA sequencing, and neuroanatomical mapping, we demonstrate that the Esr1+ LHA-LHb pathway receives a major input from a striosomal Tac1+/Tshz1+/Oprm1+ ACB neuron subtype. Intersectional cell-type-specific and input-output defined optogenetic manipulation of this ACB-LHA-LHb pathway revealed its role in signaling aversion after repeated activation, with the negative behavioral state being dependent on recruitment of Esr1+ LHA-LHb neurons. Importantly, we found that activation of the D1+ ACB-LHA pathway drives reward-independent compulsive-like seeking behaviors, expressed as compulsive digging or poking behaviors. We found that these complex yet stereotyped behaviors compete with highly motivated states and can override the need for natural rewards or social stimuli. Our findings reveal a discrete striosomal Tac1+ ACB projection targeting the aversive Esr1+ LHA-LHb pathway as a key circuit that promotes compulsive seeking behaviors over goal-directed actions.

## INTRODUCTION

Animals must flexibly adapt their behavior in response to contextual cues and homeostatic signals, balancing goal-directed actions with exploration or foraging, as well as expressing complex innate behaviors such as grooming and social interactions^1,2^. The hypothalamic circuitry plays a central role in the dynamic competition between behavioral states and actions by integrating external stimuli and internal states to support appropriate behaviors^3^. Suppression of repetitive and non-rewarded actions is essential for maintaining appropriate context-dependent responses, and impairments in the ability to flexibly adapt behaviors are central to compulsivity and stereotypies. The hypothalamic circuitry has been implicated in conditions where maladaptive and compulsive behaviors are prominent, such as obesity, anorexia, obsessive-compulsive disorder, and addiction^4^.

The hypothalamus is spatially organized and composed of a large number of neuron subtypes^5–8^. Within the hypothalamus, neurons in the lateral hypothalamic area (LHA) directly control motivational states and behavioral programs essential for maintaining energy homeostasis (e.g. feeding)^9^. GABAergic and glutamatergic neurons in the LHA have been identified as having opposing roles in regulating feeding behaviors, with GABAergic neurons promoting feeding and glutamatergic neurons inhibiting it^10^. LHA neurons that drive feeding have also been linked to compulsive and stereotyped actions, for example Agrp-expressing neurons, which were first identified as central regulators of hunger signals^11^, can additionally promote the expression of compulsive or stereotyped behaviors^12^.

The general role of LHA neurons in motivated behaviors has been established, yet it remains unknown how circuit-specific inputs and outputs of LHA neuron subtypes shape the balance and competition between behavioral states. The integration of internal homeostatic signals with value prediction signals and external stimuli depends on projection of hypothalamic signals to downstream circuits. For example, LHA glutamatergic projections to the ventral tegmental area (VTA) control reward and compulsive seeking without impacting food consumption^10^ and have a role in stress-induced overfeeding^13^. Instead, a distinct pathway of LHA glutamatergic neurons projecting to the lateral habenula (LHb), a hub for processing aversive stimuli and negative reinforcement, control negative states and feeding^14–18^. The identity and function of the glutamatergic LHA-LHb neurons are complex^19^; with one subtype expressing estrogen receptor 1 (Esr1) being central for aversive signaling and in negative stress-related states^20^. Glutamatergic neurons in the LHA receive inputs from the bed nucleus of stria terminalis (BST) that can suppress feeding and promote aversion^21^. Another major input that controls feeding behavior comes from D1+ ACB neurons that target GABAergic as well as glutamatergic LHA neurons^22,23^.

To study the circuit mechanisms in LHA that control the competition between aversive states and the expression of goal-directed actions, we investigated the organization and function of the cell-type specific inputs to the aversive LHA neuron subtype that is defined by expression of the estrogen receptor 1 (Esr1) and projections to the lateral habenula (Esr1+ LHA-LHb neurons). We mapped the identity of the whole-brain synaptic inputs to Esr1+ LHA-LHb neurons using neuroanatomical and single-neuron RNA-seq profiling and performed circuit-specific manipulation during naturalistic and motivated behaviors. In summary, our findings establish the organization and identity of a striosomal type Tac1+/Tshz1+ ACB-LHA projection that promotes reward-independent compulsive-like seeking behaviors.

## RESULTS

### Characterization of monosynaptic inputs to the Esr1+ LHA-LHb pathway

We first used a strategy to map the cell-type-specific monosynaptic inputs to the LHA neuron subtype defined by its projections to LHb and expression of estrogen receptor 1 (Esr1+ LHA-LHb neurons), using a genetically modified rabies virus for whole-brain input tracing^24,25^. We injected an adeno-associated virus (AAV) with cre-dependent expression of the TVA receptor and the rabies glycoprotein (RG) into the LHA of Esr1-cre mice followed by injection of the EnvA-coated GFP-expressing rabies virus into LHb (Figure 1a, b). Based on the GFP-labeling of the whole-brain monosynaptic inputs, we quantified the input populations following the region definitions in the Allen mouse brain reference atlas (Figure 1c). We found that the largest number of labeled inputs were found locally within the hypothalamus, while the largest number of long-range inputs were found in the cerebral nuclei (Figure 1d). We further analyzed the distribution of the labeled inputs in the cerebral nuclei and found that the largest proportion of labeled inputs was localized in the nucleus accumbens (ACB), followed by the striatum-like amygdalar nuclei (sAMY), and for the pallidum, in the bed nuclei of the stria terminalis (BST) and the substantia innominata (SI) (Figure 1e, f).

**Figure 1.**
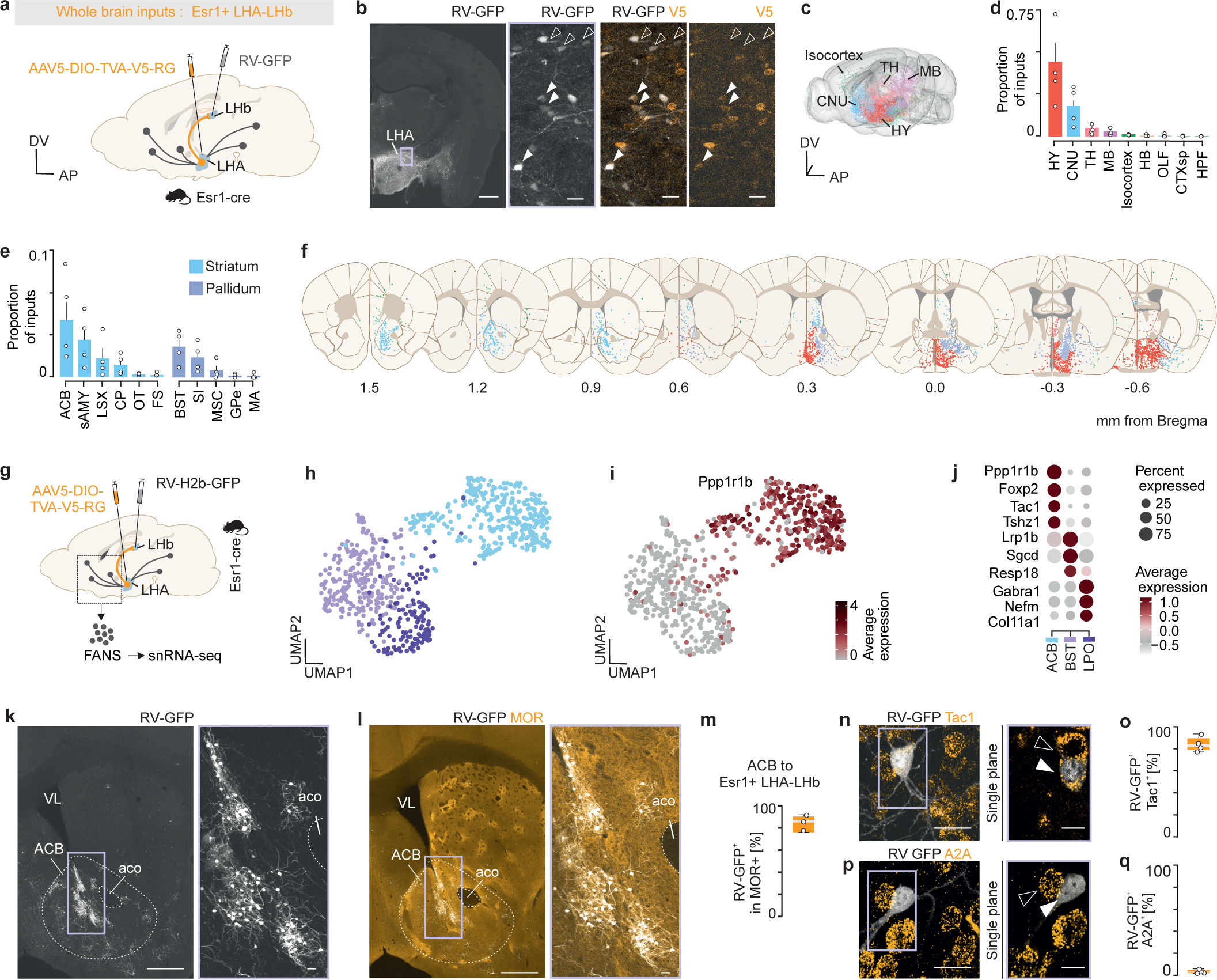
Characterization of the monosynaptic inputs to the Esr1+ LHA-LHb pathway. (a) Experimental strategy for rabies tracing of Esr1+ LHA-LHb inputs. The helper virus (AAV5-DIO-TVA-V5-RG) was injected in the LHA, and the EnvA-coated rabies virus (RV-GFP) in the LHb. (b) Representative confocal images of starter neurons co-expressing RV-GFP (white) and V5 (orange; Esr1-cre mouse; light purple box: location of right panels; arrowheads: RV-GFP+/V5+ neurons; empty arrowheads: RV-GFP+/V5-neurons). (c) Whole-brain visualization of RV-GFP+ labeled neurons (color-coded by neuroanatomical region; 21742 neurons, n = 1). (d,e) Quantification of inputs per brain region (n = 4 mice). (f) Coronal plates (Allen Brain Atlas CCFv2) showing detected GFP+ input neurons (one dot = one neuron, each plate shows neurons detected in two consecutive brain sections from one representative mouse). (g) Rabies tracing strategy and workflow for single-nucleus RNA sequencing (snRNA-seq) of inputs to Esr1+ LHA-LHb neurons. (h) UMAP plot showing clustering of neurons based on their transcriptional profile (1093 nuclei, average 4744.375 genes/neurons). (i) Visualization of the striatal marker Ppp1r1b in the UMAP plot. (j) Average expression of selected cell type markers per cluster. (k) Representative confocal images of ACB inputs to Esr1+ LHA-LHb (GFP in white; light purple box: location of right panel). (l) Same image as in (k), showing mu-opioid receptor (MOR) expression (orange; light purple box: location of right panel). (m) RV-GFP+ ACB neurons overlapping with MOR expression. (n) Representative confocal images showing Tac1 expression (orange) in ACB RV-GFP+ neurons (white; light purple box: location of right panel, arrowheads: RV-GFP+/Tac1+ neurons; empty arrowheads: RV-GFP-/Tac1+ neurons). (o) Quantification of Tac1 expression in RV-GFP+ACB neurons. (p) Representative confocal images showing A2A expression (orange) in ACB RV-GFP+ neurons (white; light purple box: location of right panel, arrowheads: RV-GFP+/A2A+ neurons; empty arrowheads: RV-GFP-/A2A+ neurons). (q) Quantification of A2A expression in RV-GFP+ACB neurons. Scale bars: 500 μm (b left, k left, l left), 20 μm (b right, k right, l right, n, p). For boxplots (m,o,q), data shown as median (centerline), box (25th and 75th percentiles), whiskers (non-outlier minimum and maximum) and outliers (>1.5 interquartile range). Abbreviations: anterior commissure olfactory limb (aco), lateral ventricle (VL), fluorescence-activated nuclear sorting (FANS), cerebral nuclei (CNU), thalamus (TH), hypothalamus (HY), midbrain (MB), olfactory areas (OLF), cerebral cortex cortical subplate (CTXsp), hippocampal formation (HPF), nucleus accumbens (ACB), striatum-like amygdala (sAMY), lateral septal complex (LSX), caudoputamen (CP), olfactory tubercle (OT), fundus of striatum (FS), bed nuclei of the stria terminalis (BST), substantia innominata (SI), medial septal complex (MSC), Globus pallidus external segment (GPe), magnocellular nucleus (MA).

To identify the molecular profile of neurons subtypes with inputs to Esr1+ LHA-LHb neurons, we labeled the monosynaptic inputs using a nuclear-localized GFP (H2b-GFP) expressing rabies virus followed by single nuclei isolation and single-nucleus RNA sequencing (snRNA-seq) (Figure 1g). Analysis of the snRNA-seq data revealed that the input populations clustered into three main groups, with the largest cluster consisting of Ppp1r1b+ neurons (Figures 1h, i). The Ppp1r1b+ cluster confirmed that the major input to Esr1+ LHA-LHb neurons came from striatal projection neurons, and we therefore further analyzed cell type markers in the striatal cluster to identify possible ACB subtypes. We found that ACB neurons with monosynaptic inputs to Esr1+ LHA-LHb neurons were defined by expression of several striatal neuron subtype markers (e.g., Foxp2, Tac1, Tshz1; Figure 1j), that supported the notion that ACB projections to Esr1+ LHA-LHb neurons are formed by a discrete Tac1+/Tshz1+ pathway.

Based on the expression of the striosome-enriched markers Tac1 and Tshz1 in the ACB subpopulation, we imaged the spatial organization of the rabies-labeled neurons, which revealed a characteristic striosomal type organization, with GFP+ labeled neurons being co-localized with the dense mu opioid receptor (MOR) staining (Figure 1k-m). To further profile the GFP+ labeled ACB neurons, we performed in situ hybridization for the two main neuron types in ACB, namely the D1 and D2-receptor expressing neurons, using Tac1 and A2A expression as cell type markers. We found that 85% of the GFP+ labeled ACB neurons could be classified as Tac1+ neurons (Figure 1n-q). In summary, using brain-wide monosynaptic rabies tracing together with snRNA-seq and neuron type profiling we identified that the Esr1+ LHA-LHb neurons receive a major input from Tac1+/Tshz1+/Oprm1+ striosomal type ACB neurons.

### Neuroanatomical mapping of a cell-type-specific ACB-LHA-LHb circuit

To directly visualize the distribution of the striosomal type ACB synaptic terminals in hypothalamus, we injected an AAV enabling cre-dependent expression of tdTomato together with a synaptophysin-localized GFP fusion protein (SYP-GFP) into the ACB of D1-cre or Oprm1-cre mice (Figure 2a-c). We found for both D1+ as well as Oprm1+ ACB neurons, the localization of SYP-GFP expressing axon terminals concentrated to a specific domain in the LHA which is consistent with the canonical location of the Esr1+ LHA-LHb population^20^ (Figure 2d-g). To further confirm the cell-type specific organization of the LHA-LHb neurons receiving ACB inputs, we designed an intersectional input-output somatic mapping strategy to genetically label all neurons that receive inputs from ACB and that also project to LHb. We used an anterograde transsynaptic mapping strategy based on AAV1 injection (AAV1-Cre) into the ACB to label the postsynaptic targets of the ACB neurons, in combination with labeling of the LHb-projecting neurons using injections of retrogradely transported cre-dependent AAV (AAVretro-DIO-tdTomato) into the LHb (Figure 2h). Our intersectional input-output mapping strategy resulted in the genetic labeling specifically of the LHA-LHb neurons that receive direct ACB inputs (Figure 2i). This labeling strategy showed that the LHA-LHb subpopulation was topographically distributed in a LHA domain that overlaps with the densest axonal projections from D1+ and Oprm1+ ACB neurons, and the majority of the labeled neurons expressed Esr1 (Figure 2j-l).

**Figure 2.**
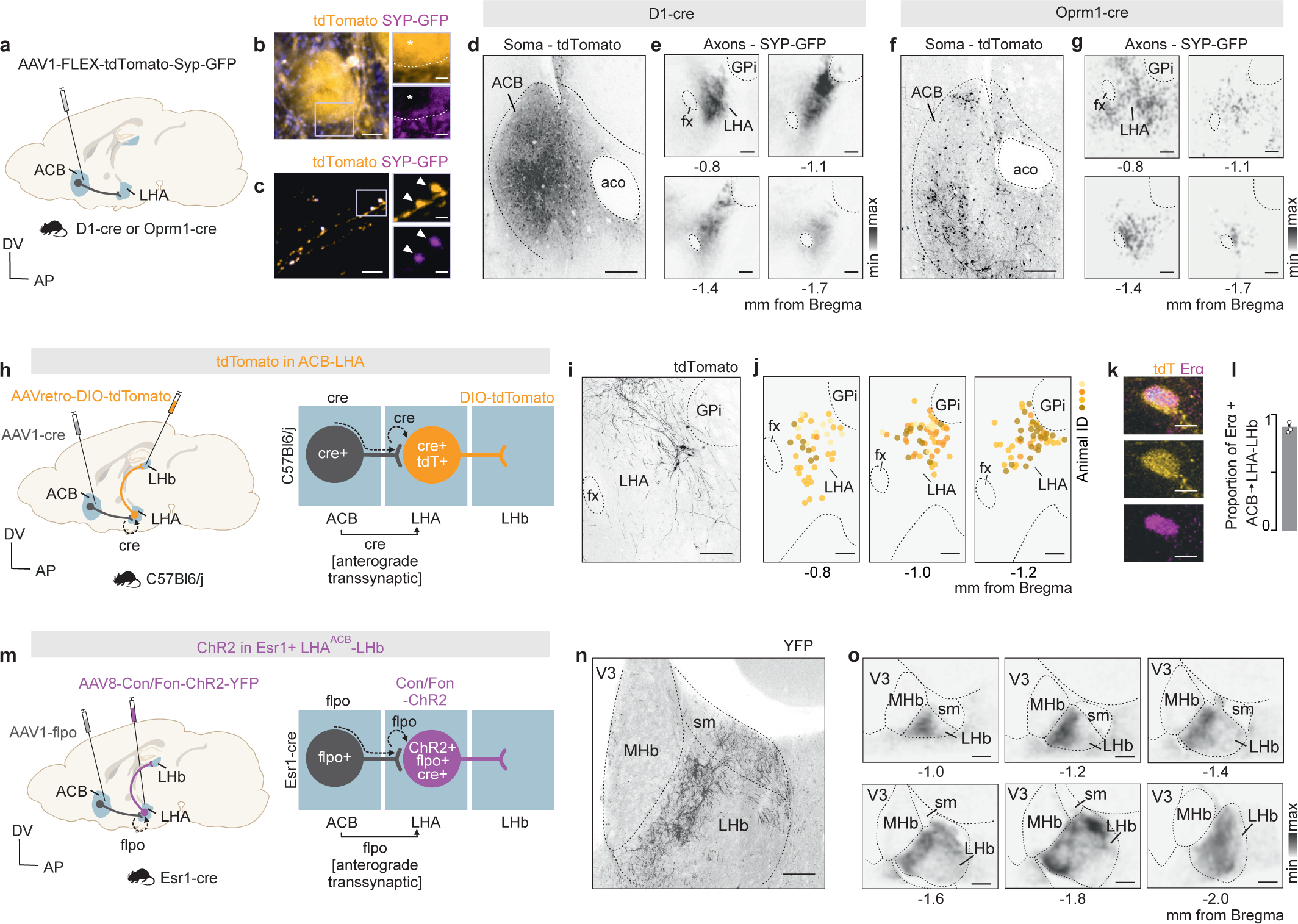
Neuroanatomical mapping of a cell-type-specific ACB-LHA-LHb circuit. (a) Viral strategy for cell-type specific labeling of ACB-LHA axon terminals. (b-c) Representative confocal images showing expression of tdTomato (orange) in soma (b, soma in ACB) and axons (c, axons in LHA), and of SYP-GFP to presynaptic terminals (purple; light purple boxes: location of right panels; asterisk: tdTomato+ soma; arrowheads: ACB-LHA tdTomato+ /SYP-GFP+ presynaptic terminal). (d) Representative confocal image of tdTomato expression in ACB D1+ neurons. (e) Heatmaps of ACB D1+ axon terminal distribution in LHA (black, SYP-GFP+). (f) Representative confocal image of tdTomato expression in ACB Oprm1+ neurons. (g) Heatmaps of ACB Oprm1+ axon terminal distribution in LHA (black, SYP-GFP+). (h) Experimental strategy for ACB-input and LHb-output specific labelling of LHA neurons. (i) Representative confocal image of transsynaptic labeling of LHA-LHb neurons with ACB inputs. (j) Coronal plates showing the position of detected LHA-LHb neurons with ACB inputs (n= 4 mice; one color = one mouse; one dot = one neuron). (k) Representative confocal image showing Erα protein expression in LHA-LHb neurons with ACB inputs (tdTomato, orange; Erα, purple). (l) Quantification of Erα expression in LHA-LHb neurons with ACB inputs. (m) Experimental strategy transsynaptic labeling of Esr1+ LHA-LHb neurons with ACB inputs and expression of ChR2. (n) Representative confocal image showing labeling of axon terminals in LHb from Esr1+ LHA-LHb neurons with ACB inputs. (o) Heatmaps of axon terminal distribution in LHb from Esr1+ LHA neurons with ACB inputs (black, SYP-GFP+). Scale bars: 300 μm (d, f), 250 μm (i), 100 μm (e, g, j, n, o), 10 μm (k), 2 μm (b left, c left), 1 μm (b right, c right). Abbreviations: anterior commissure olfactory limb (aco), Fornix (fx), Globus pallidus internus (GPi), medial habenula (MHb), stria medullaris (sm), third ventricle (V3).

To directly visualize the axonal projection in LHb of Esr1+ LHA neurons that receive inputs from ACB, we used a different input-output axonal mapping strategy. We injected transsynaptic AAV1-flpo into the ACB and a cre-dependent/flpo-dependent vector for expression of ChR2 (AAV8-Con/Fon-hChR2-YFP) into the LHA of Esr1-cre mice (Figure 2m). This strategy resulted in the genetic expression of ChR2-YFP in Esr1+ LHA neurons with specific inputs from ACB, and visualization of their axon terminals in LHb (Figure 2n). We found that Esr1+ LHA-LHb neurons receiving ACB inputs showed a spatially organized axonal targeting within the LHb, with a preferential targeting of the medial domain of the LHb (Figure 2o). In summary, we established intersectional input-output mapping strategies to target Esr1+ LHA-LHb neurons receiving ACB inputs and mapped their organization in LHA as well as their discrete target domain in the LHb.

### Repeated activation of D1+ ACB-LHA projection neurons drive aversion through the Esr1+ LHA-LHb pathway

To investigate how modulation of aversive states depends on the integration of the ACB inputs on the Esr1+ LHA-LHb pathway, we first tested the aversive signaling of LHA-LHb neurons that are genetically defined by Esr1+ expression and ACB inputs. To specifically target the Esr1+ LHA-LHb neurons with direct inputs from ACB, we used an intersectional input-output strategy in Esr1-cre mice, based on transsynaptic anterograde AAV1 labeling to express flpo recombinase only in neurons that receive ACB inputs, together with injection into the LHA of the flpo-dependent and cre-dependent ChR2 vector (AAV8-Con/Fon-ChR2-YFP). With this intersectional strategy we expressed ChR2 only in Esr1+ LHA-LHb neurons targeted by the ACB, and we placed optical fibers on top of LHb to perform axon terminal stimulation (Figure 3a). We used a real-time place avoidance test to quantify the ChR2-induced aversive signals, with a first exposure (10 min) of optogenetic activation in one compartment, followed by a second exposure (10 min) where the optogenetic activation was switched to the opposite compartment (Figure 3b). We found that mice strongly avoided the compartment with the ChR2-induced activation of the ACB-targeted Esr1+ LHA-LHb terminals in the first exposure as well as after switching compartment stimulation (Figure 3c). This result confirmed our prediction that Esr1+ LHA-LHb neurons with specific ACB inputs drive strongly aversive signals.

**Figure 3.**
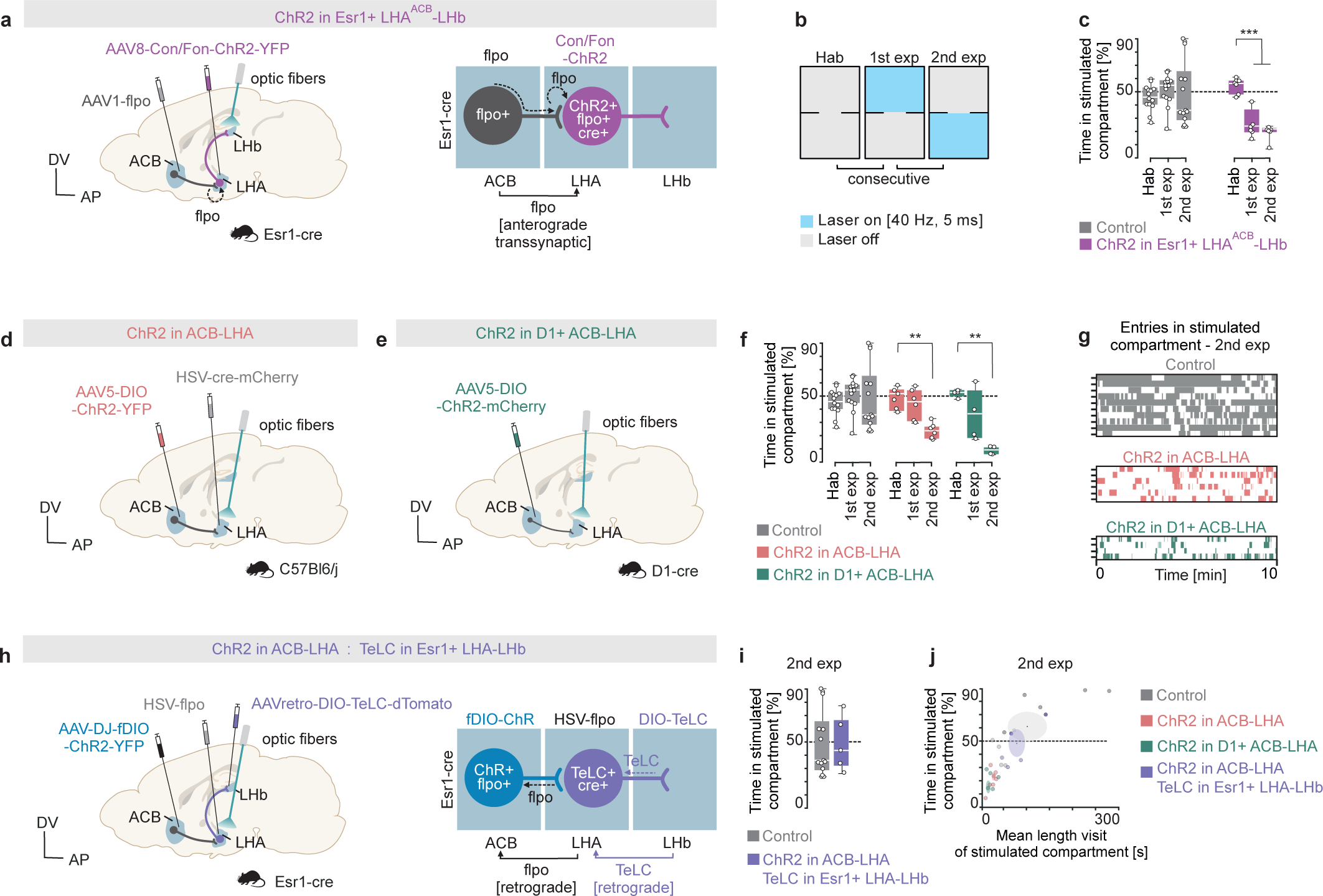
Repeated activation of D1+ ACB-LHA projection neurons drive aversion through the Esr1+ LHA-LHb pathway. (a) Experimental strategy for optogenetic terminal stimulation of Esr1+ LHA-LHb neurons with direct ACB inputs (ChR2 in Esr1+ LHA^ACB^-LHb). (b) Schematic of the real-time place avoidance test. (c) Optogenetic activation of Esr1+ LHA^ACB^-LHb pathway decreased time spent in the stimulated compartment during first and second exposure of the real-time place avoidance test (RM two-way ANOVA, group x test phase: F(2,32) = 6,146, p = 0.0055; ChR2 in Esr1+ LHA^ACB^-LHb Hab vs 1st exp: p = 0.0006, ChR2 in Esr1+ LHA^ACB^-LHb Hab vs 2nd exp: p < 0.0001, Holm-Sidak’s post-hoc test; control n = 14; ChR2 in Esr1+ LHA^ACB-^LHb, n = 7). (d) Strategy for pathway-specific optogenetic activation of ACB-LHA axon terminals (ChR2 in ACB-LHA). (e) Strategy for cell-type-specific optogenetic activation of D1+ ACB-LHA axon terminals (ChR2 in D1+ ACB-LHA). (f) Optogenetic activation of D1+ ACB-LHA or ACB-LHA pathways induced an avoidance of the stimulated compartment during the second exposure in the real-time place avoidance test (RM two-way ANOVA, group x test phase: F(4,36) = 3.462, p = 0.0171; ChR2 in ACB-LHA Hab vs 2nd exp: p = 0.0095; ChR2 in D1+ ACB-LHA Hab versus 2nd exp: p = 0.0012, Holm-Sidak’s post-hoc test; control n = 14; ChR2 in ACB-LHA n = 6; ChR2 in D1+ ACB-LHA n = 4). (g) Visualization of entries to stimulated compartment (one color coded vertical bar = 1 sec in stimulated compartment). (h) Experimental strategy for optogenetic activation of ACB-LHA pathway combined with TeLC silencing of the Esr1+ LHA-LHb neurons (ChR2 in ACB-LHA: TeLC in Esr1+ LHA-LHb). (i-j) Optogenetic activation of ACB-LHA pathway together with TeLC silencing of Esr1+ LHA-LHb pathway normalized the mean duration and number of visits to the stimulated compartment (control n = 14; ChR2 in ACB-LHA and TeLC in Esr1+ LHA-LHb n = 5; ChR2 in ACB-LHA n = 6; ChR2 in D1+ ACB-LHA n = 7). For boxplots (c,f,i), data shown as median (centerline), box (25th and 75th percentiles), whiskers (non-outlier minimum and maximum) and outliers (>1.5 interquartile range). * p < 0.05, ** p < 0.01, *** p < 0.001.

To assess the function of ACB inputs to LHA in modulating aversive states, we used two different viral strategies for optogenetic activation of ACB terminals that target the LHA as well as the cell-type-specific activation of the D1+ ACB-LHA projection neurons (Figure 3d, e). In the first strategy, we injected the retrogradely transported Herpes simplex virus (HSV) into LHA to express cre recombinase (HSV-cre-mCherry) in neurons projecting to LHA and a cre-dependent ChR2-expressing vector into ACB (AAV5-DIO-ChR2-YFP), resulting in a selective targeting of LHA projecting ACB neurons. In the second strategy, we targeted D1+ projection neurons in ACB using local injection of a cre-dependent ChR2-expressing vector in D1-cre mice (AAV5-DIO-ChR2-mCherry) and stimulated their axon terminals through optical fibers placement in the LHA. In contrast to the immediate avoidance induced by direct stimulation of the Esr1+ LHA-LHb pathway, we found that activation of the inhibitory ACB inputs in LHA did not immediately induce preference or avoidance for the ChR2-stimulated compartment in the real-time place avoidance test. Interestingly, in response to the second exposure and repeated ChR2-stimulation we found an increased and significant avoidance to the ChR2-stimulated compartment (Figure 3f, g). These results together suggest that activation of the D1+ ACB terminals in LHA does not produce an immediate place preference through inhibition of the LHA-LHb pathway, but after repeated activation instead induces an aversive state.

We therefore asked if the negative state induced by repeated optogenetic activation of ACB-LHA neurons depended on recruitment of the Esr1+ LHA-LHb pathway. To test this hypothesis, we used an intersectional input-output strategy that allowed us to perform optogenetic stimulation of the ACB-LHA projections together with cell-type specific silencing of Esr1+ LHA-LHb neurons using tetanus light chain (TeLC) expression. We targeted the ACB neurons that project to the LHA using an HSV-based retrograde expression of flpo (HSV-flpo in LHA) combined with a local injection of the flpo-dependent ChR2 in ACB (AAV-DJ-fDIO-ChR2-YFP) in Esr1-cre mice. We combined this projection-specific ChR2 expression with cell-type specific silencing of the Esr1+ LHA-LHb neurons using injection into LHb of a retrograde vector with cre-dependent TeLC expression (AAVretro-DIO-TeLC-dTomato) (Figure 3h). We found that the cell-type-specific silencing of the Esr1+ LHA-LHb neurons completely blocked the avoidance response of repeated activation of ACB-LHA projections (Figure 3i, j). In summary, we found that the D1+ ACB-LHA pathway shapes the transition to an aversive state after repeated activation, and that this negative state depends on recruitment of the Esr1+ LHA-LHb neurons.

### The D1+ ACB-LHA pathway drives a compulsive-like digging state

Following our findings on the negative state induction after repeated activation of the ACB-LHA pathway in the place avoidance assay, we aimed to define how activation impacted the expression of discrete exploratory or motor motifs in an open field environment. To capture the detailed behavioral changes imposed by activation of the D1+ ACB-LHA pathway, we used automated tracking and pose estimation using DeepLabCut (DLC^26^) and classified behavioral motifs using an unsupervised probabilistic deep learning model (VAME^27^) (Figure 4a). We found that repeated optogenetic activation of D1+ ACB-LHA terminals resulted in a significant increase in locomotor speed and shifted the behavioral repertoire towards vigorous exploratory motifs such as rearing and fast turning (e.g. motifs defined as “fast walk and stand”, “fast walk and sharp turn”; Figure 4b-e). We then asked whether these behavioral motifs represented a general negative state or stress-related response that would generalize to an enriched home-cage environment. Therefore, we again optogenetically activated the D1+ ACB-LHA axon terminals in an enriched environment consisting of natural bedding material and freely available food (pellets) (Figure 4f). Unexpectedly, in this context we found that mice developed a strong digging behavior upon activation of the D1+ ACB-LHA axon terminals (Figure 4g).

**Figure 4.**
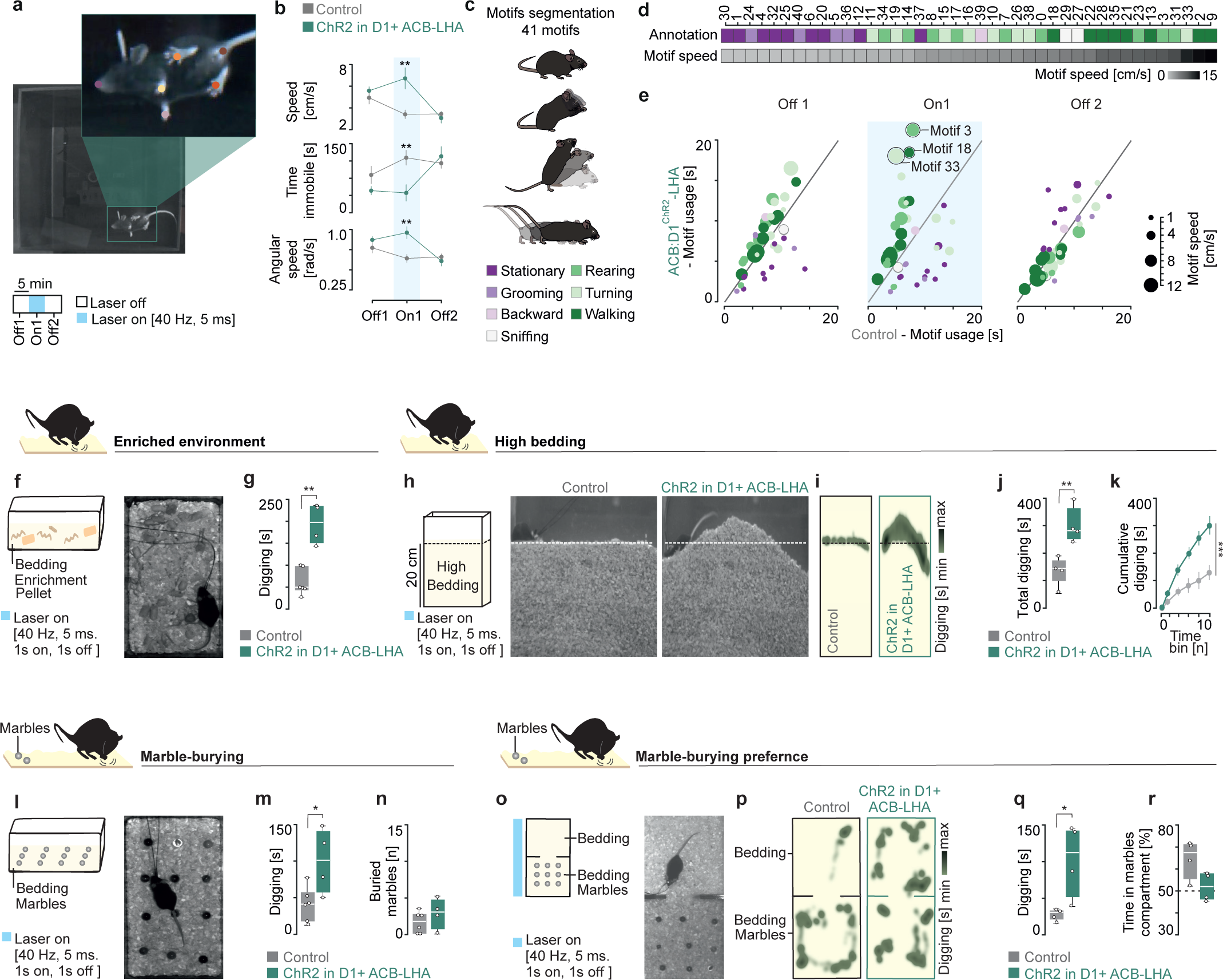
The D1+ ACB-LHA pathway drives a compulsive-like digging state. (a) Schematic of the open field test. (b) Optogenetic activation of D1+ ACB-LHA pathway increased locomotor and angular speed while decreasing immobility during open field test (Control vs ChR2 in D1+ ACB-LHA, speed: p On1 = 0.008, time immobile: p On1 = 0.007, angular speed: p On1 = 0.003; two-way ANOVA with Bonferroni’s post-hoc test; control, n = 14; ChR2 in D1+ ACB-LHA, n = 10). (c) Schematic of motifs segmented using Variational Autoencoders for Motif Extraction (VAME). (d) Behavioral annotations and relative speed of the 41 motifs found with VAME. (e) Visualization of motif usage during optogenetic activation of D1+ ACB-LHA pathway vs control. Each motif is represented with a point (color = behavioral annotation, size = motif speed; motif 3: fast walk and stand; motif 18: straight walk; motif 33 = walk and sharp turn). Same mice as in (b). (f) Schematic and image of the enriched home cage assay. (g) Optogenetic activation of the D1+ ACB-LHA pathway increased time spent digging in the enriched home cage assay (Mann-Whitney, control vs ChR2 in D1+ ACB-LHA: p = 0.0095; control, n = 6; ChR2 in D1+ ACB-LHA, n = 4). (h) Schematic and image of the high bedding assay. (i) Heatmaps of the digging location in the arena. (j,k) Optogenetic activation of D1+ ACB-LHA pathway increased digging behavior across the entire session (Mann-Whitney, total digging control vs ChR2 in D1+ ACB-LHA: p = 0.0317) (RM two-way ANOVA, group x time bin: F(5,35) = 8,258, p < 0.0001; control, n = 4; ChR2 in D1+ ACB-LHA, n = 4). (l) Schematic and image of the marble burying test. (m,n) Optogenetic activation of D1+ ACB-LHA pathway increased digging behavior without changing the number of buried marbles (Mann-Whitney, digging control vs ChR2 in D1+ ACB-LHA, p = 0.0242; control, n = 6; ChR2 in D1+ ACB-LHA, n = 4). (o) Schematic and image of the two-compartments marble-burying assay. (p) Heatmaps of the digging location in the two compartments of the arena. (q,r) Optogenetic activation of ACB-LHA pathway increased digging behavior without changing the time spent in marble-paired compartment (Mann-Whitney, digging control vs ChR2 in D1+ ACB-LHA, p = 0.0317; control, n = 4; ChR2 in D1+ ACB-LHA, n = 4. For boxplots (f,j,m,n,q,r), data shown as median (centerline), box (25th and 75th percentiles), whiskers (non-outlier minimum and maximum) and outliers (>1.5 interquartile range). * p < 0.05, ** p < 0.01, *** p < 0.001.

We therefore performed the same ACB-LHA activation paradigm in a context designed to promote exploratory and digging behavior using a high bedding assay (Figure 4h). In this context, we found that mice showed a strong and sustained digging behavior during the entire period of optogenetic activation of the D1+ ACB-LHA axons (Figure 4h-k). We next asked if the sustained digging behavior represented a goal-directed or purposeful action or was the expression of a stress-related and compulsive-like response. For this, we performed D1+ ACB-LHA terminal activation in the marble-burying test and found a significant increase in the digging behavior during optogenetic stimulation, but this response was surprisingly not associated with increased marble burying (Figure 4l-n). To test for a potential preference between digging and marble-burying, we performed a modified place-preference test where bedding material was present in both compartments while marbles were paired with only one compartment (Figure 4o). Optogenetic stimulation of D1+ ACB-LHA terminals induced significant digging behavior irrespective of the marble-paired compartment, and no preference for the marble-paired compartment (Figure 4p-r). In summary, we found that activation of a discrete D1+ ACB-LHA pathway leads to sustained and compulsive-like seeking or digging behavior that is not directed towards salient objects (e.g. marbles).

### The D1+ ACB-LHA-LHb pathway drives context-independent compulsive-like seeking behaviors

The induction of a persistent and compulsive-like digging behavior during activation of the ACB-LHA projections motivated us to further define the context-dependent nature of the behavioral response. We first used a hole-board assay to study exploration, seeking, and poking behaviors in a novel environment (Figure 5a). We found that mice with optogenetic activation of the D1+ ACB-LHA axon terminals showed a significant and sustained increase in hole poking in the absence of salient stimuli or natural rewards (e.g. food) (Figure 5b). To directly test the role of the downstream LHA-LHb neurons in mediating the compulsive-like hole poking, we established an intersectional input-output strategy to optogenetically silence with NpHR specifically the LHA-LHb neurons that receive ACB inputs (Figure 5c). This strategy resulted in a significant and sustained increase in hole poking during the hole-board assay (Figure 5d-f) supporting that inhibition of the LHA-LHb pathway mediates the compulsive-like poking behavior induced by activation of the D1+ ACB-LHA pathway.

**Figure 5.**
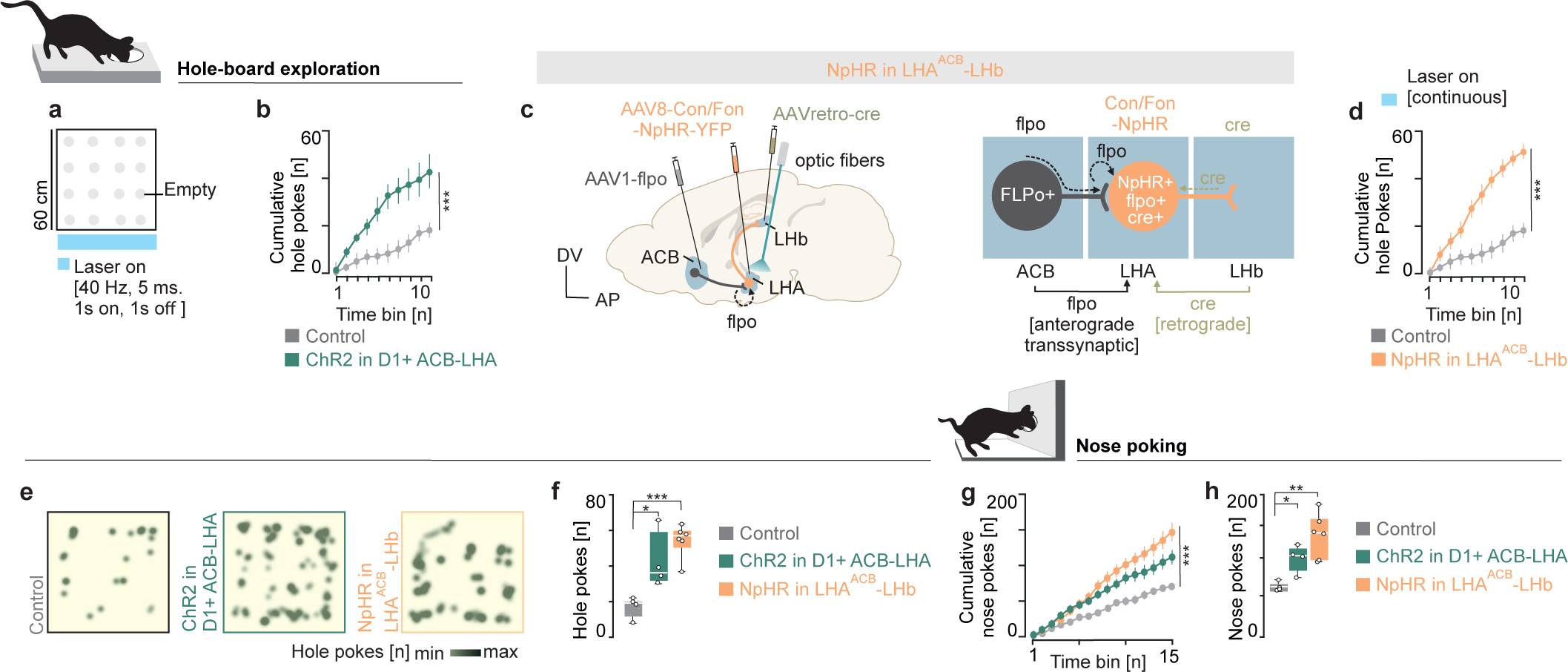
The D1+ ACB-LHA-LHb pathway drives context-independent compulsive-like seeking behaviors. (a) Schematic of the hole-board assay. (b) Optogenetic activation of D1+ ACB-LHA pathway increased hole-poking behavior (RM two-way ANOVA, group x time bin: F(9,54) = 7.043, p < 0.0001; control, n = 4; ChR2 in D1+ ACB-LHA, n = 4). (c) Intersectional strategy for optogenetic silencing of LHA-LHb neurons receiving ACB inputs (NpHR in LHA^ACB^-LHb). (d) Optogenetic silencing of the LHA-LHb pathway receiving ACB inputs increased hole-poking behavior (RM two-way ANOVA, group x time bin: F(9,72) = 24.54, p < 0.0001). Control, n = 4; NpHR in LHA^ACB^-LHb, n = 6. (e) Heatmaps of hole-poking in the hole-board assay. (f) Optogenetic activation of D1+ ACB-LHA pathway or optogenetic silencing of LHA-LHb pathway receiving ACB inputs increased hole-poking (unpaired one-way ANOVA: F(2,14) = 10.99, p = 0.014; control vs ChR2 in D1+ ACB-LHA: p = 0.012; control vs NpHR in LHA^ACB^-LHb: p = 0.0007, Holm-Sidak’s post-hoc test; control, n = 4; ChR2 in D1+ ACB-LHA, n = 4; NpHR in LHA^ACB^-LHb, n = 6). (g,h) Optogenetic activation of D1+ ACB-LHA pathway as well as optogenetic silencing of LHA-LHb neurons with ACB inputs increased the number of non-rewarded nose pokes across the entire session (RM two-way ANOVA, group x time bin: F(30,165) = 5.255, p < 0.0001; Cumulative nose pokes control vs ChR2 in D1+ ACB-LHA: p < 0.0001; Cumulative nose pokes control vs NpHR in LHA^ACB^-LHb: p < 0.0001, Holm-Sidak’s post-hoc test; unpaired one-way ANOVA: F(2,11) = 11.22, p = 0.0022; total nose pokes control vs ChR2 in D1+ ACB-LHA: p = 0.04; total nose pokes control vs NpHR in LHA^ACB^-LHb: p = 0.0012, Holm-Sidak’s post-hoc test). Same mice as in (f). For boxplots (f,h), data shown as median (centerline), box (25th and 75th percentiles), whiskers (non-outlier minimum and maximum) and outliers (>1.5 interquartile range). * p < 0.05, ** p < 0.01, *** p < 0.001.

To investigate whether the compulsive-like poking is a form of generalized seeking behavior that can be expressed in other contexts, we tested how D1+ ACB-LHA activation impacted unrewarded nose-poking behavior. In this assay, we found that the two circuit manipulation strategies, either optogenetic activation of D1+ ACB-LHA axon terminals or silencing of LHA-LHb neurons receiving ACB inputs, both resulted in a persistent and significant increase in nose-poking behavior in the absence of any natural reinforcer (Figure 5g-h). Together, the results from the two different manipulation strategies strongly support the central role of a D1+ ACB projection to LHA that drives reward-independent compulsive-like seeking behavior through inhibition of the LHA-LHb pathway.

### The D1+ ACB-LHA pathway suppresses competing motivational states to drive compulsive-like seeking

We further aimed to define how competing high-motivational states could impact the expression of the compulsive-like seeking behavior. We therefore activated the D1+ ACB-LHA pathway in water-deprived mice that were trained to nose-poke for a liquid reward (water drop) in a progressive ratio paradigm. We found that mice with D1+ ACB-LHA pathway activation showed a nose-poking behavior that was motivation and reward-independent. Supporting the motivation-independent nature of the poking behavior, mice showed constant rate of nose-poking over the entire session that resulted in an overall similar number of total pokes and total consumed rewards as the highly-motivated control group (Figure 6a-d). These results suggest that the compulsive-like behaviors persisted even in the presence of natural rewards and competing high-motivation states.

**Figure 6.**
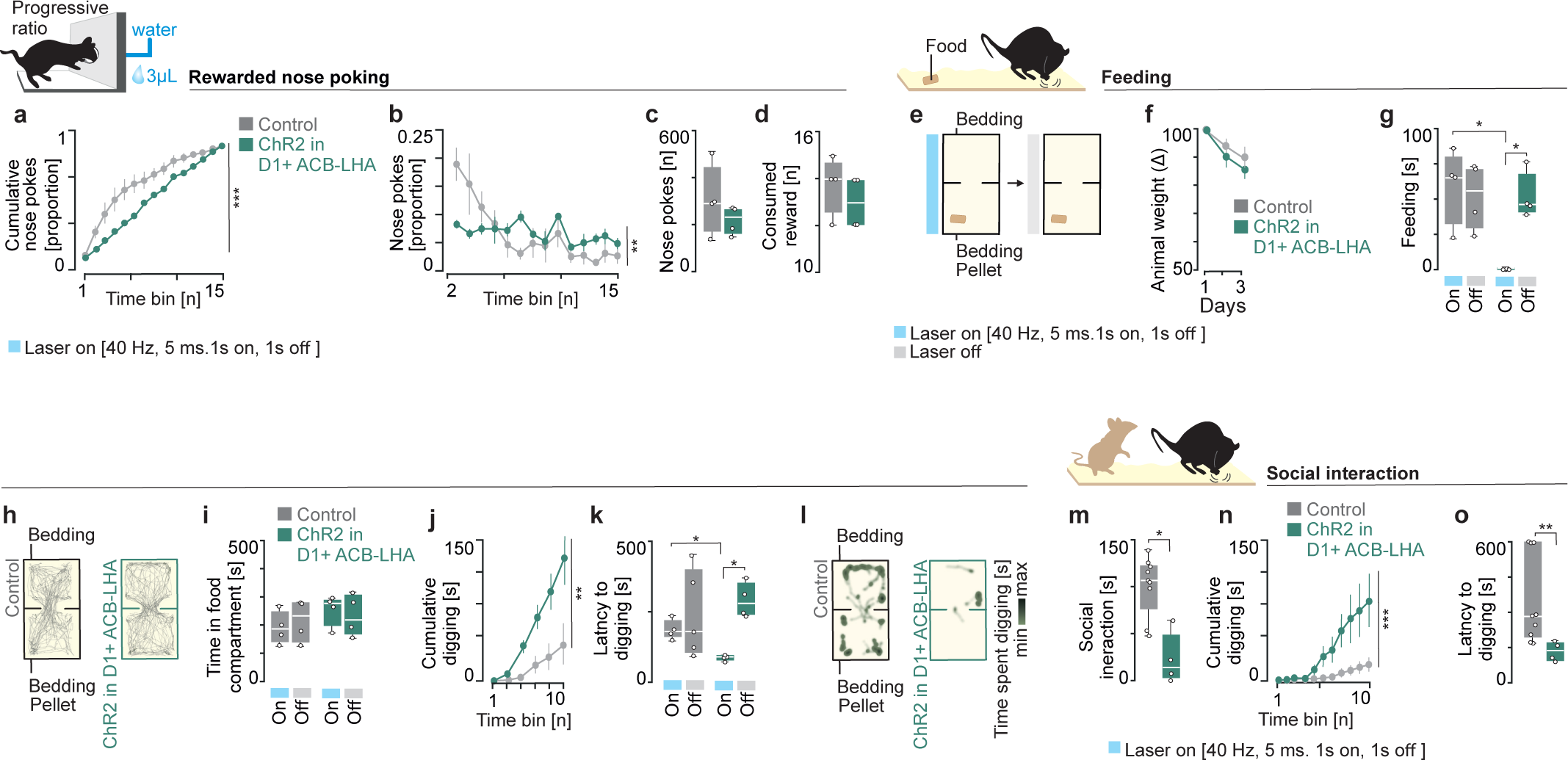
The D1+ ACB-LHA pathway suppresses competing motivational states to drive compulsive-like seeking. (a,b) Optogenetic activation of D1+ ACB-LHA pathway in water-restricted mice induced a constant reward-independent nose-poking frequency in the progressive-ratio test (RM two-way ANOVA, group x time bin: F(14,84) = 5.393, p < 0.0001) (RM two-way ANOVA, group x time bin: F(13,78) = 2.897, p = 0.0018). control, n = 4; ChR2 in D1+ ACB-LHA, n = 4. (c,d) Optogenetic activation of D1+ ACB-LHA pathway did not change the amount of nose-poking or consumed rewards. Same mice as in (a,b). (e) Schematic of the two-compartment food test. (f) Weight reduction in food-restricted mice (control, n= 4; ChR2 in D1+ ACB-LHA, n = 4). (g) Optogenetic activation of D1+ ACB-LHA pathway suppressed feeding behavior (Mann-Whitney, control On vs ChR2 in D1+ ACB-LHA On: p = 0.0286; ChR2 in D1+ ACB-LHA On vs ChR2 in D1+ ACB-LHA Off: p = 0.0286). Same mice as in (a-f). (h) Example of mouse tracking in the two-compartment food test. (i) Optogenetic activation of D1+ ACB-LHA pathway did not change the time spent in the food-paired compartment. Same mice as in (a-g). (j) Optogenetic activation of D1+ ACB-LHA pathway increased digging behavior across the entire session in the two-compartment food test (RM two-way ANOVA, group x time bin: F(5,30) = 5.191, p = 0.0015). (k) Optogenetic activation of D1+ ACB-LHA pathway decreased the latency to first digging behavior in the two-compartment food test (Mann-Whitney, control On vs ChR2 in D1+ ACB-LHA On: p = 0.0286; ChR2 in D1+ ACB-LHA On vs ChR2 in D1+ ACB-LHA Off: p = 0.0286). Same mice as in (a-j). (l) Heatmaps showing digging in the two-compartment food test. (m) Optogenetic activation of D1+ ACB-LHA pathway decreased social interaction time (Mann-Whitney, control vs ChR2 in D1+ ACB-LHA: p = 0.0112). Same mice as in (a-k). (n) Optogenetic activation of D1+ ACB-LHA pathway increased digging behavior across the entire session (RM two-way ANOVA, group x time bin: F(10,110) = 9.12, p < 0.0001). Same mice as in (a-k). (o) Optogenetic activation of D1+ ACB-LHA pathway induced short-latency digging behavior in the social interaction test (Mann-Whitney, control vs ChR2 in D1+ ACB-LHA: p = 0.0098). For boxplots (c,d,g,i,k,m,o), data shown as median (centerline), box (25th and 75th percentiles), whiskers (non-outlier minimum and maximum) and outliers (>1.5 interquartile range). * p < 0.05, ** p < 0.01, *** p< 0.001.

Next, to determine whether the compulsive-like behaviors could override other motivational states, we activated the D1+ ACB-LHA pathway in a feeding and in a social interaction context. To determine the competition with feeding programs, we activated the D1+ ACB-LHA pathway in food-deprived mice in a modified two-compartment assay with bedding in both compartments and freely-accessible food (pellet) in only one compartment (Figure 6e, f). In this assay, hungry mice with optogenetic activation of the D1+ ACB-LHA pathway showed a complete suppression of feeding behavior. Notably, feeding behavior was restored when the optogenetic activation ended (Figure 6g). Importantly, the reduced feeding behavior was not coupled to avoidance of the food-paired compartment (Figure 6h, i). Instead, we found that the D1+ ACB-LHA activation induced a significant rapid increase in digging behavior in both compartments (Figure 6j-l). Finally, we assessed whether the observed prioritization of compulsive-seeking behavior over consummatory actions (e.g., water or food) could also override other competing motivational states. To test this, we exposed mice to an unfamiliar, sex-matched mouse and in a home-cage environment with bedding and quantified the social interaction. In this context, activation of the D1+ ACB-LHA pathway shifted the behavior towards compulsive-like digging over social interaction (Figure 6m-o). Altogether, these findings uncover a central role for the D1+ ACB-LHA pathway in promoting and prioritizing compulsive stereotyped behaviors, even in the presence of competing high-motivation states such as thirst, hunger, and social interaction.

## DISCUSSION

Hypothalamic circuits are known to control homeostatic and innate behaviors, and to play a role in motivated behaviors. The motivation to engage in specific actions is influenced by rewarding as well as aversive signals, with the lateral hypothalamic (LHA) to habenula (LHb) projection having a central function in valence processing. More specifically, glutamatergic projections from LHA that target the LHb induce strong aversion and depression-like behaviors^16,18^, mediated by a neuroanatomically, molecularly, and physiologically distinct Esr1+ population in LHA that shows sex-specific sensitivity in the development of stress-related states^20^.

Here we describe a molecularly and neuroanatomically discrete striosomal Tac1+/Oprm1+ subtype in ACB that targets Esr1+ LHA-LHb neurons. Based on the connectivity profile, we hypothesized that activation of the GABAergic ACB-LHA projection would produce an immediate positive signal through direct inhibition of the Esr1+ LHA-LHb neurons, leading for example to preference for a compartment associated with optogenetic stimulation of the D1+ ACB-LHA terminals. Surprisingly, we found that activation of the ACB-LHA pathway did not induce an immediate preference response, and instead over repeated stimulation induced a negative or avoidance state. We interpreted the absence of an immediate behavioral effect after activation of the D1+ ACB-LHA pathway and the subsequent avoidance upon repeated stimulation as a network effect or plasticity mechanism to support integration of circuit activity over longer timescales for the execution of complex behavioral programs. Surprisingly, we found that the D1+ ACB-LHA projection drives complex yet stereotyped compulsive-like digging and nose-poking behaviors even in the presence of natural rewards or social stimuli, indicating a shift in behavioral priority. This shift suggests that the activation of the ACB-LHA-LHb circuit can override physiological motivational hierarchies, and upon repeated activation promote compulsive-like behavioral programs over goal-directed actions. The maladaptive network or plasticity effects of such circuit activation overlap with the concept of habit formation^28^ and the transition from goal-directed to compulsive behaviors in addiction models^29^.

The role of LHA in reinforcing behaviors has been revealed through electrical self-stimulation experiments^30,31^, and electrical stimulation experiments have neuroanatomically mapped in LHA subregions the induction of stereotyped or compulsive-like behaviors such as grooming and digging^32^. Supporting the complex role of hypothalamic circuits structuring and selecting between behavioral programs, it has for example been shown that Agrp-expressing neurons and feeding overrides other motivational programs^33^, while Agrp-expressing neurons have also been implicated in driving stereotypic actions like foraging and digging, behaviors that can be associated with food seeking or energy deficits^12^. In addition, other hypothalamic circuits (e.g. LHA projections to PVH) control the balance between competing behaviors, such as feeding and stress-induced compulsive-like behaviors (e.g. grooming)^34^, and galanin-expressing LHA neurons reduces compulsive marble burying whereas activation of all GABAergic LHA neurons increases this behavior^35^. The temporal sequence of LHA-mediated behavioral repertoires shows some underlying structure, where some behavioral states tend to precede others, for example in the case of nest-building before sleep that has been linked to LHA activity, including LHA neurons with projections to LHb^36^. These findings in summary highlight the broader and more complex function of hypothalamic feeding-related circuits, also offering insights into conditions defined by repetitive behaviors, such as obsessive-compulsive disorders. The connection between feeding circuits and repetitive or stress-related behaviors like marble-burying, further underscores the importance of hypothalamic networks in maintaining behavioral and physiological homeostasis.

A key circuit in mediating motivation, reward, and compulsivity is the nucleus accumbens, a key component of the mesolimbic reward system^37^. ACB neurons are diverse in terms of their molecular profile and projection pattern^38,39^, with D1+ striatal projection neurons generally associated with promoting motivated behaviors, while D2+ neurons linked to aversion and inhibitory control^40,41^. Interestingly, our results reveal that D1+ ACB neurons provide a monosynaptic input to Esr1+ LHA-LHb neurons and challenge the current view where D1+ neurons discreetly promote reward-related behaviors. This agrees with evidence that D1+ neuron subtypes in ACB can encode negative motivational valence^42,43^ and spatially organized Pdyn+/D1+ subtypes drive either aversion or reward^44^. Our findings support a more complex view on the diverse function of ACB neurons, by demonstrating that D1+ ACB neuron subtypes with circuit-defined downstream targets can shape behaviors beyond reinforcement.

In summary, we found a striosomal Tac1+/Tshz1+/Oprm1+ monosynaptic input from ACB to Esr1+ LHA-LHb neurons that drives reward-independent compulsive-like seeking behaviors, even in the presence of strongly competing motivational states such as feeding and social interaction. Our work contributes to a circuit-based understanding of the transition between competing behavioral states, which can uncover the underlying mechanisms of psychiatric conditions where compulsive behaviors are prominent.

## METHODS

### Animals

All procedures and animal experiments were performed according to the guidelines of the Stockholm Municipal Committee for animal experiments and the Karolinska Institutet in Sweden (approval number 155440-2020). Adult mice aged 3–8 months were used: Esr1-cre (B6N.129S6(Cg)-Esr1tm1.1(cre)And/J; Jackson Laboratory Stock, 017911), D1-cre (B6.FVB(Cg)-Tg(Drd1-cre)EY262Gsat/Mmucd; MMRRC Stock, 030989-UCD), Oprm1-ce (generated in Meletis lab^45^), and wild-type mice (C57BL/6J; Charles River). Animals were group housed in a temperature (23 °C) and humidity (55%) controlled environment in standard cages on a 12:12 h light/dark cycle with ad libitum access to food and water, unless placed on a food/water restriction schedule. All food/water restricted mice were restricted to 85–90% of their initial body weight by administering one feeding of 2.0–2.5 g of standard grain-based chow per day for food restriction and 1 mL water for water restriction. All strains used were backcrossed with the C57BL/6J strain. Male and female mice, when possible aged matched and littermates, were used for viral tracing experiments, in vivo optogenetic manipulation and behavioral testing. The number of animals (n) for each experiment is reported in the corresponding figure legend.

### Tracers and viral constructs

Purified and concentrated AAVs were purchased from Addgene: AAV1-phSyn1-FLEX-tdTomato-T2A-SypEGFP (51509-AAV1), AAV1-hSyn-Cre (105553-AAV1), AAVrg-Ef1α-fDIO-mCherry (114471-AAVrg), AAV1-EF1α-Flpo (55637-AAV1), pAAV-hSyn Con/Fon hChR2(H134R)-EYFP (55645-AAV8), pAAV-EF1α-DIO-hChR2(H134R)-EYFP (20298-AAV5), pAAV-EF1α-DIO-hChR2(H134R)-mCherry (20297-AAV5), pAAV-EF1α-Cre (55636-AAVrg), and pAAV-nEF-Con/Fon-NpHR3.3-EYFP (137152-AAV8). HSV-EF1α-mCherry-IRES-Cre and HSV-EF1α-Flpo were obtained from the Viral Gene Transfer Core, MIT. ssAAV-DJ/2-shortCAG-dFRT-hChR2-EYFP (v237-DJ) and ssAAV-retro/2-hSyn1-dlox-TETxLC_2A_NLS_dTomato (v620-retro) were purchased from VVF Zurich. For rabies tracing, helper virus AAV-EF1α-DIO-TVA-V5 and EnvA-coated rabies viruses (RV-GFP and RV-H2B-GFP) were produced in the Meletis lab.

### Stereotaxic injections

Stereotaxic injections were performed on 8–12-week-old mice. Mice were anesthetized with isoflurane (3% for induction, 1–2% for maintenance). Body temperature was maintained at 37°C using a heating pad, and ocular ointment (Viscotears, Alcon) was applied to protect the eyes. The head was fixed in a stereotaxic apparatus (Kopf), and lidocaine (4 mg/kg) was injected locally before the skin incision. After cleaning the skull with chlorhexidine, craniotomies (∼300–500 μm) were performed. Unless otherwise specified, all injections were carried out at a speed of 100 nl/min using a glass micropipette connected to a Quintessential Stereotaxic Injector (Stoelting). Viral labeling of ACB-LHA neurons was achieved by bilaterally injecting 200 nl of anterograde cre-dependent synaptophysin-expressing virus (AAV1-phSyn1(S)-FLEX-tdTomato-T2A-SypEGFP-WPRE) into the ACB. The ACB was targeted using the following coordinates: 1.1 mm rostral, 0.65 mm lateral to bregma, and 3.7 mm deep from the dura.

For the neuroanatomical mapping of the ACB-LHA-LHb pathway, 200 nl of an anterograde transsynaptic Cre-expressing virus (AAV1-hSyn-Cre-WPRE-hGH) was injected into the ACB, and retrograde flpo-dependent mCherry-expressing virus (AAVretro-Ef1α-fDIO-mCherry) was injected into the LHb at a rate of 100 nl/min. This injection was performed unilaterally, with the LHb targeted using the coordinates: 1.65 mm caudal, 0.4 mm lateral to bregma, and 2.55 mm deep from the dura.

To visualize Esr1+ specific ACB-LHA-LHb axons, 300 nl of anterograde transsynaptic flpo-expressing virus (AAV1-EF1α-Flpo) was injected into the ACB, and 300 nl of cre and flpo dependent YFP-expressing virus (AVV5-Con/Fon-ChR2-eYFP) was injected into the LHA. This injection was done unilaterally, and the LHA was targeted using the following coordinates: 1.25 mm caudal, 1 mm lateral to bregma, and 4.65 mm deep from the dura.

For optogenetic stimulation of the ACB-LHA-LHb pathway, a bilateral injection of 250 nl of anterograde transsynaptic flpo-expressing virus (AAV1-EF1α-Flpo) into the ACB and 250 nl of cre and flpo dependent YFP-expressing virus (AAV8-Con/Fon-ChR2-eYFP) into the LHA was performed at a rate of 80 nl/min. To stimulate the ACB-LHA pathway, two different approaches were employed: the first involved a bilateral injection of 200 nl of local cre-dependent ChR2-expressing virus (AAV5-EF1α-DIO-hChR2(H134R)-EYFP-WPRE-HGHpA) into the ACB and 200 nl of retrograde cre-expressing virus (HSV-EF1α-mCherry-IRES-Cre) into the LHA. The second approach used a bilateral injection of 200 nl of cre-dependent ChR2-expressing virus (AAV5-EF1α-DIO-hChR2(H134R)-mCherry-WPRE-HGHpA) into the ACB.

For stimulation of the ACB-LHA pathway in combination with silencing of the Esr1 LHA-LHb pathway, we established an intersectional approach using bilateral injection of 200 nl of flpo-dependent ChR2-expressing virus (AAVDJ-FRT-chR2-EYFP) into the ACB, 200 nl of retrograde flpo-expressing virus (HSV-EF1α-flpo) into the LHA, and 200 nl of retrograde cre-dependent tetanus-expressing virus (ssAAVretro/2-hSyn1-dlox-TETxLC_2A_NLS_dTomato(rev)-dlox-WPRE-hGHp(A)) into the LHb. Another approach to silence the ACB-projecting LHA-LHb pathway involved a bilateral injection of 250 nl of retrograde cre-expressing virus (AAVretro-EF1α-Cre) into the LHb, along with 250 nl of anterograde transsynaptic flpo-expressing virus (AAV1-EF1α-Flpo) into the ACB and 250 nl of cre and flp dependent NpHR3.3-expressing virus (AAV8-nEF-Con/Fon-NpHR3.3-EYFP) into the LHA at a rate of 50 nl/min. Optogenetic experiments were conducted 15 days after injection. TeLC experiments were performed 4 weeks post-injection.

For the rabies tracing experiment, 150 nl of helper virus (AAV-EF1α-DIO-TVA-V5-WRPE-hGHpA) was first injected into the LHA, followed three weeks later by injection of 150 nl of RV-GFP into the LHb at a rate of 80 nl/min. For the snRNA-seq experiment, 200 nl of the same helper virus was injected into the LHA followed three weeks later by injection of 200 nl RV-H2B-GFP into the LHb (injection rate 50 nl/min). The mice were euthanized 10 days after the injection of the RV-GFP.

For anatomical studies, mice were euthanized 3 weeks post-injection. In all stereotaxic injections, the glass pipette was left in place for 10 minutes post-injection before being removed from the brain. Buprenorphine (0.1 mg/kg i.p.) was administered after anesthesia, before surgery and carprofen (5 mg/kg i.p.) was administered immediately after surgery and 24 hours later for pain relief.

### Implant surgery

Mice were anesthetized with isoflurane (3% for induction, 1–2% for maintenance). Body temperature was maintained at 37°C using a heating pad, and ocular ointment (Viscotears, Alcon) was applied to protect the eyes. The head was fixed in a stereotaxic apparatus (Kopf), and lidocaine (4 mg/kg) was injected locally before the skin incision. After cleaning the skull with chlorhexidine, two small craniotomies (∼300–500 μm) were made to insert the fibers, which were lowered vertically into the brain for photoactivation (LHb: +1.65 mm AP, 0.95 mm ML, 2.2 mm depth, 10° angle; LHA: +1.1 mm AP, 1.2 mm ML, 4 mm depth). The fibers were secured with light-curing dental adhesive (OptiBond FL, Kerr) and cement (Tetric EvoFlow, Ivoclar Vivadent). Buprenorphine (0.1 mg/kg i.p.) was administered after anesthesia, before surgery and carprofen (5 mg/kg i.p.) was administered immediately after surgery and 24 hours later for pain relief.

### Immunohistochemistry

Brain sections were cut on a vibratome at 50 μm thickness (Leica VT1000, Leica Microsystems GmbH). For rabies tracing and Esr1 staining of the ACB-LHA-LHb pathway, the sections were rinsed in phosphate buffer (PB), blocked for 2 hours in 10% normal donkey serum with 0.5% Triton X-100, and then incubated overnight at 4°C with primary antibodies. The primary antibodies used were chicken anti-V5 (1:1000; Abcam, ab9113) and rabbit anti-ESRα (1:5000; Millipore, 06-935.) Following extensive PB washing, sections were incubated using Cy3-and Cy5-conjugated secondary antibodies (1:1000; Jackson ImmunoResearch, 703-165-155 and 711-175-152, respectively) for 2 hours at room temperature. All antibodies were diluted in a carrier solution consisting of PB with 1% bovine serum albumin (BSA), 1% normal goat serum, and 0.5% Triton X-100. After further rinsing in PB, the sections were mounted on slides and coverslipped.

For MOR staining sections were rinsed in TBST (TBS with 0.3% Triton), blocked for 1 hour in 10% donkey serum in TBS at room temperature with gentle agitation, and incubated overnight with the primary antibody rabbit anti-MOR (1:500; Abcam, ab134054) diluted in 2% donkey serum in TBST at room temperature. After washing with TBST, the sections were incubated for 2 hours at room temperature with a Cy5-conjugated secondary antibody (1:1000, same as above) diluted in 2% donkey serum in TBST. Following extensive washing with TBST and PBS, the sections were mounted on slides and coverslipped.

### RNAscope

The RNAscope Fluorescent Multiplex Assay V2 kit was used on rabies-GFP-expressing brain sections mounted on microscope slides. Samples were washed in 1X PBS, then treated with Pretreat 2 solution at 70-80°C for 5 minutes. After washing with distilled water and 100% ethanol, Protease3 (1 drop) was added and incubated for 30 minutes at 40°C. Following another wash, 1 drop of the appropriate probe (Tac1 or A2A) was applied and incubated for 2 hours at 40°C. The sections were washed with 1X wash buffer, and 1 drop of AMP-1FL was added, followed by 30 minutes of incubation at 40°C. This procedure was repeated for AMP-2FL (15 minutes) and AMP-3FL (30 minutes). After washing with 1X wash buffer, HRP-C1 was added and incubated for 15 minutes at 40°C. Cy5 TSA-fluorophore (1:500) was applied and incubated for 5 minutes at 40°C, followed by GRP-blocker for 15 minutes at the same temperature. Finally, after a wash with 1X wash buffer, DAPI (1:1000) was applied, and the sections were coverslipped.

### Single-nucleus RNA sequencing

To isolate brain tissue, mice were first perfused with 15mL of the following solution: 80 mM NaCl, 75 mM sucrose, cutting/recovery solution (10x), 20 mM D-glucose (10x), 26 mM NaHCO3 (10x). 300 µm sections were cut using a vibratome and immediately put in a RNALater solution. The main input regions were isolated and kept in -20 °C until processing for nuclei extraction. To extract nuclei, tissue was transferred into a 2 mL tube with NP40 Lysis Buffer (prepared according to the Nuclei Isolation from Complex Tissues for Single Cell Multiome ATAC + Gene Expression Sequencing protocol, CG000375) and got homogenized with the use of 2 step homogenizer. After incubation of 5 minutes in NP40 Lysis Buffer, the homogenate was filtered through a 40 µm strainer and transferred into microcentrifuge tube to be centrifuged with 500 x g for 5 minutes at 4 °C. After supernatant was removed, the pellet was incubated in PBS with 1% BSA and 1 U/µL RNA inhibitor, and then resuspended. Then, this step was repeated with the same centrifuge settings and solutions. 1:1000 DAPI was added to the final solution, and it was sorted in a FACS machine for the GFP signal. The collection tubes were coated with BSA beforehand. Then the nuclei were fixed using the Parse Evercode Nuclei Fixation v3 kit. ∼735 nuclei were taken as input for Parse Evercode WT Evercode v3 Library preparation (using 3 wells of the kit). In situ nuclei barcoding, cDNA capture and amplification, and sequencing library preparation were performed according to the manufacturer’s instructions and samples were split into 8 sublibraries after in situ nuclei barcoding. For both cDNA amplification and Index PCR, 8 cycles of PCR were performed respectively. The length distribution profile of the final library was confirmed with the Agilent Bioanalyzer, using the high-sensitivity DNA chip, yielding an average fragment size of 463 bp. Sublibraries were pooled to a final concentration of 19.56 ng/uL, and samples were sent to Xpress Genomics AB for circularisation, DNA Nanoball making, and sequencing on a MGI DNBSEQT7 Machine (PE150 kit) using Illumina TruSeq sequencing primers. The read layout was set up according to the Parse Biosciences instructions (read1: 146bp, read2: 58bp, index1: 8bp, index2: 8bp) with stable linker sequences in the Parse barcode region being dark cycle sequenced. FASTQ files were demultiplexed into the respective sublibraries and preprocessed using the ParseBiosciences-Pipeline (v1.3.0), accounting for dark cycled bases by adding the flag “bc_amp_seq NNNNNNNNNN333333332222222211111111” to the parfile. After pre-processing with default parameters of the Parse pipeline, 1093 cells passed initial QC (that is, containing a full and unique parse barcode sequence). Reads were mapped to the mouse genome (mm39) using the internal version of STAR of the parse pipeline. Transcripts were annotated using gencode transcript annotations (vM31).

### Image acquisition

Confocal images were captured using a Zeiss 880 confocal microscope and exported via ZEN black (2.1 SP3 v14.0). For viral expression overview, a Plan-Apochromat 10×/0.45 M27 objective was used with the following settings: frame size 1,024 × 1,024; pinhole 1.59 AU; bit depth 16-bit; speed 7; averaging 2. For detailed imaging of viral expression and immunohistochemistry, a Plan-Apochromat 20×/0.8 M27 objective was employed with settings: frame size 1,024 × 1,024; pinhole 1.07 AU; bit depth 8-bit; speed 6; averaging 2. For in situ RNA quantification, a Plan-Apochromat 63×/1.40 Oil DIC M27 objective was used with settings: frame size 1,024 × 1,024; pinhole 0.85 AU; bit depth 8-bit; speed 6; averaging 4.

### Optogenetics

Mice were bilaterally implanted with optical fibers targeting the LHb (coordinates: +1.65 mm AP, 0.95 mm ML from bregma, and 2.2 mm depth at a 10° angle from the dura) or the LHA (coordinates: +1.1 mm AP, 1.2 mm ML from bregma, and 4 mm depth from the dura). The implanted optical fibers were purchased from RWD Life Science (R-FOC-BL200C-22NA, 3 mm for LHb, 5 mm for LHA). Mice were connected via a splitter branching patch cord (SBP(2)_200/220/900-0.22_1m_FCM-2xMF1.25, Doric Lenses) to their implanted optical fibers using a split sleeve (ADAL1-5, Thorlabs). The splitter branching patch cord was connected to a laser (MLL-III-447-200mW laser) via a fiber-optic rotary joint (FRJ_1×1_FC-FC, Doric Lenses) to prevent cable twisting during animal’s movement. The frequency and duration of photostimulation were controlled using a custom-written Arduino script (Arduino IDE), through Bonsai software (v2.6.3). For optogenetic stimulation, light power was measured in continuous light at 8 mW and for optogenetic inhibition power was measured in continuous light at 2 mW at the tip of the splitter branching patch cord cable before each experiment using an optical sensor (Thorlabs).

### Behavioral tests

#### Real-time place avoidance test

Mice were placed in a custom-made two-compartment behavioral arena, constructed from black plexiglass and measuring 50 × 25 × 25 cm, separated by a wall with an opening in the middle. The behavioral arena was situated on a transparent plexiglass surface, and animal behavior was recorded using a camera positioned below the arena. During habituation the optical fibers were connected to the animal without light stimulation. During the first exposure one compartment was paired with light stimulation (40 Hz, 5 ms pulse, 447 nm laser, 10 min) followed by a second exposure in the opposite compartment (10 min). Mice were tracked using Ethovision XT 17, and the time spent in each compartment was recorded.

#### Open field test

Mice were placed in a custom-made open field (30 x 30 cm) made of black plexiglass for 15 minutes. The arena was positioned on a transparent plexiglass surface to allow the camera to record behavior from below at 20 fps. Mouse performance was evaluated during alternating epochs of laser stimulation, beginning with a 5-minute off period, followed by 5-minute stimulation epochs (40 Hz, 5 ms pulse, 1s on-1s off, 447 nm laser). Body point pose estimation and motif segmentation in the videos were performed using DeepLabCut (DLC) and Variational Autoencoders for Motif Extraction (VAME).

#### Enriched home cage assay

The experiment began with the mouse being placed in an empty clean cage (30 x 15 cm), where an optogenetic stimulation was applied for 5 minutes (40 Hz, 5 ms pulse, 1s on-1s off period, 447 nm laser; not recorded). While keeping the optogenetic stimulation ongoing, the mouse was then placed back in its home cage and recorded for 10 minutes. Mice were tracked using Ethovision XT 17, and digging behavior was manually scored.

#### High bedding assay

In a glass arena measuring 40 x 40 x 6 cm, 20 cm of sawdust bedding was placed. The experiment began by placing the mouse in a clean, empty cage for 5 minutes of optogenetic stimulation (40 Hz, 5 ms pulse, 1s on-1s off, 447 nm laser; not recorded). Following this, the mouse was placed in the arena, where it was recorded for 10 minutes while stimulation continued. Mouse movements were tracked using Ethovision XT 17, and digging behavior was manually scored.

#### Marble burying test

12 clean glass marbles, each with a diameter of 1.5 cm and a homogeneous color, were placed in a clean cage containing 2.5 cm thick layer of bedding made from sawdust, without food or water. The experiment began with the mouse being placed in an empty clean cage, where optogenetic stimulation was applied for 5 minutes (40 Hz, 5 ms pulse, 1s on-1s off period, 447 nm laser; not recorded). After this initial stimulation, the mouse was placed in the cage with the marbles for 10 minutes while the stimulation continued. At the end of this period, each mouse was returned to its cage, and the number of buried marbles was scored; marbles were considered buried if they were covered by sawdust by at least two-thirds. Before testing a new mouse, the marbles were cleaned with 70% ethanol and placed in a newly prepared cage. Mice were tracked using Ethovision XT 17, and discrete events of digging were manually scored for each second.

#### Two-compartment marble-burying assay

In a custom-made two-compartment behavioral arena (50 × 25 × 25 cm), one black and one white compartment were separated by a wall with a central opening. Both compartments contained a 2.5 cm layer of sawdust bedding, with no food or water present. Nine clean glass marbles (1.5 cm diameter) were placed in the black compartment. The experiment began with the mouse in a clean cage, receiving optogenetic stimulation for 5 minutes (40 Hz, 5 ms pulse, 1s on-1s off, 447 nm laser; not recorded). After this, while stimulation continued, the mouse was placed in the arena and recorded for 10 minutes. At the end of the session, each mouse was returned to its cage, and the number of buried marbles was counted, with marbles considered buried if covered by sawdust by at least two-thirds. Before testing a new mouse, marbles were cleaned with 70% ethanol and placed in a freshly prepared cage. Mice were tracked using Ethovision XT 17, recording the time spent in each compartment and manually scoring discrete digging events.

#### Hole-board assay

Mice were food restricted overnight. The experiment began with the mouse being placed in an empty clean cage, where optogenetic stimulation was applied for 5 minutes (40 Hz, 5 ms pulse, 1s on-1s off period, 447 nm laser; not recorded). Following this, the mouse was placed in an arena 45 x 45 cm featuring 16 empty holes, to allow exploration of the apparatus during the first 10 minutes while continuing optogenetic stimulation. The holes were situated on a heightened platform, enabling hole poking. Mice were tracked using Ethovision XT 17, and discrete events of poking in the holes were manually scored.

#### Nose-poking test (unrewarded)

Mice were placed in a 15 x 15 cm box with one nose-poke port. Water delivery tubing was removed, and no reward was given. The experiment began by placing the mouse in a clean, empty cage with optogenetic stimulation (5min, 40 Hz, 5 ms pulse, 1s on-1s off, 447 nm laser). The mouse was then placed in the nose-poking box, where the number of pokes was recorded for 15 minutes.

#### Progressive-ratio test

Mice were water-restricted (85-90% of their initial body weight) and received 1 mL of water per day. Mice were trained for 4 days without optogenetic stimulation while connected to a patch cord cable. Training proceeded as follows: Day 1 - one reward per nose poke; Day 2 - one reward per two nose pokes; Days 3 and 4 - one reward per five nose pokes. On Day 5, mice underwent 5 minutes of stimulation in a clean cage before a 15-minute progressive ratio test, where one reward was given for an increasing number of nose pokes, doubling the required pokes after every two rewards (i.e., 1, 1, 2, 2, 4, 4, 8, 8, 16, 16, 32, 32, 64, 64, 128, 128).

#### Two-compartment food test

Mice were food-restricted to 85 to 90% of their initial body weight by administering one daily feeding of approximately 2.5 to 3.0 g of standard grain-based chow, which was given immediately following the behavioral experiment. The mice underwent food restriction for 48 hours prior to the experiment, and their weight was measured. The experiment began with the mouse being placed in an empty clean cage, where optogenetic stimulation was applied for 5 minutes (40 Hz, 5 ms pulse, 1s on-1s off period, 447 nm laser; not recorded). Following this, food-restricted mice were placed in a custom-made two-compartment arena, consisting of a black compartment and a white compartment, separated by a wall with an opening in the middle (50 × 25 × 25 cm). Both compartments contained a 2.5cm thick layer of bedding to enable digging behavior. One food pellet was placed in the white compartment, and the mouse was allowed to freely consume it for 10 minutes while the optogenetic stimulation continued. Mice were recorded using a camera positioned above the arena and were tracked with Ethovision XT 17. Discrete events of digging and eating were manually scored.

#### Social interaction test

Mice were single-housed for 90 minutes prior to the test in a cage with bedding from their home-cage. The experiment began by placing the mouse in a clean, empty cage for 5 minutes of optogenetic stimulation (40 Hz, 5 ms pulse, 1s on-1s off, 447 nm laser; not recorded). The experimental mouse returned to its home cage, where a sex-matched 8 weeks old mouse was introduced. Videos were recorded for 10 minutes and analyzed using Ethovision XT 17. Discrete events of digging and social interaction were manually scored.

### Quantification and statistical analysis

#### Statistics and reproducibility

No statistical methods were used to predetermine sample sizes, and sample sizes were similar to published literature. Statistical hypothesis testing was conducted at significance level of 0.05. All mice were randomly assigned to different groups. Data collection and analysis were not performed blind to the conditions of the experiments, unless specified. All behavioral experiments were controlled by automated computer systems/scripts (for example, ARDUINO). Mice with incorrect viral targeting or misplaced optical fiber(s) were excluded. In single-cell RNA sequencing, nuclei with less than 400 unique molecular identifiers (UMIs), or more than 100000 UMIs, or fewer than 300 genes were excluded. The exact number of animals (n) for each experiment is reported in the corresponding figure legend.

#### Clustering of cell types in snRNA-seq

Data analysis for snRNA-seq was performed in R (v4.3.2) using the Seurat package (v5.1.0). Single nuclei were extracted from both hemispheres in five mice, resulting in 860 nuclei before filtering. The snRNA-seq data were generated using the Read10X_h5() function from Seurat. Genes related to ribosomal, mitochondrial, and X and Y chromosomes were removed, and the count matrix was filtered based on reads, genes, and peaks. Low-quality nuclei and genes were filtered separately for each sample using a custom function. Doublets were removed with the DoubletFinder package (v2.0.3) using a doublet score cutoff of 0.6. RNA-seq data were normalized with SCTransform using V2 regularization, regressing out cell cycle effects based on cell cycle genes in Seurat. ATAC-seq data were normalized using TF-IDF with default settings. Harmony (group.by.vars = “sample”) was applied to remove batch effects and integrate data from different samples. Cells from both RNA-seq datasets were clustered using Weighted Nearest Neighbor (WNN) analysis via the FindMultiModalNeighbors() function (parameters: reduction.list = list(’SCT_CC_harmony’), dims.list = list(1:50), k.nn = 50). Uniform Manifold Approximation and Projection (UMAP) was used to visualize the clusters (FindClusters() with Resolution = 0.7). Marker genes were used to annotate clusters, including known markers for glial cells (Microglia, Astrocytes, Oligodendrocytes, and Oligodendrocyte precursors) and neuronal types (Medium Spiny Neurons, Interneurons, and Cortical Neurons). Specific genes analyzed included Ppp1r1b, Tac1, Foxp2, Lrp1b, Sgcd, Resp18, Gabra1, Kcnab3, and Nefm.

#### Neuroanatomical analysis

We used the WholeBrain package in R (https://github.com/tractatus/wholebrain) to register confocal images to the Allen reference atlas (Allen Brain Atlas CCFv2) and for segmentation of labeled neurons. Proportions of GFP-expressing neurons in brain areas were represented as mean ± s.e.m., with individual animal data point. Starter neurons co-expressing GFP and the V5 tag were identified using immunohistochemistry with a primary antibody against V5. For MOR immunohistochemistry analysis, confocal images of sections were acquired maintaining consistent settings across animals. For annotation of MOR+ striosomal compartments, regions of interest (ROIs) were automatically detected around MOR+ dense patches in z-stacks confocal images (Imaris). We used Imaris software to quantify the number of rabies expressing neurons that co-localized Tac1 and A2A expression. Data are represented as box plots, with single animal data points.

For axon density of ACB D1 and ACB Oprm1 terminals in LHA, confocal microscope images of SYP-GFP presynaptic terminals in LHA were acquired. Acquisition settings were kept consistent across. For each brain section, image binarization, by thresholding fluorescence level and minimum area, and pixel segmentation were performed using a custom-written script in Python. Napari brainreg plugin was used to register images into the Allen reference atlas (allen_mouse_10um, Allen Brain Atlas CCFv2). A Python custom-written script was used to combine the registration of brain sections with the pixel segmentation. As a result, pixel density heatmaps of the axon terminals at different anteroposterior coordinates were generated. Each heatmap represents the sum across 300 µm in the anterior-posterior axes. The same strategy was adopted to visualize in LHb the density of ChR2-YFP axon terminals from Esr1+ LHA neurons with inputs from ACB. For the ACB-LHA-LHb pathway, the number of soma in the LHA and the proportion of colocalization with the Esr1a staining was quantified manually. Napari brainreg plugin was used to register sample images into the Allen reference atlas (Allen Brain Atlas CCFv2). Mapping of soma position was performed using a custom-written script in Python.

#### Behavioral analysis

For the open filed analysis, we used pose estimation and motif segmentation using DeepLabCut (DLC) and Variational Autoencoders for Motif Extraction (VAME). Six virtual markers were placed on key body parts (nose, tail base, paws) across 350 uniformly sampled frames from 15 videos. A residual neural network (ResNet-50) was trained to track these markers, achieving a mean error of 2.02 pixels during training and 2.49 pixels during testing. For behavioral motif identification, the unsupervised deep learning framework VAME was trained on 85 videos (1,530,000 frames) with a test fraction of 0.1, a time window of 30 frames, and a latent dimensionality of 30, converging at 50 epochs. Clustering analysis with 100 discrete states and a 1% motif usage threshold identified 41 relevant behavioral motifs within the dataset. For all other behavioral tests, analysis was performed using Ethovision XT 17. Discrete behaviors such as digging, eating, poking, and social interactions were manually scored. Data were presented as mean ± s.e.m. or in box plots, with individual animal data point.

## Acknowledgments

We thank Yang Xuan for technical assistance with rabies virus production, Alexander Wolthon for management of the mouse colony, Jonas Frisén for providing access to the FACS infrastructure. This research was supported by the Swedish Research Council (Vetenskapsrådet MH, 2018-00608 to K.M. and 2023-01919 to D.C.), the Wenner-Gren Foundation (postdoctoral funding to T.C.), Karolinska Institutet, the neuroscience strategic research area (StratNeuro) at Karolinska Institutet (postdoctoral funding to D.C.), the Swedish Brain Foundation (Hjärnfonden, grant to K.M.). We gratefully acknowledge using the core facilities at Karolinska Institutet, specifically the animal behavior core facility (ABCF), and the Biomedicum imaging facility (BIC). We acknowledge the use of the following virus constructs: AAV phSyn1(S)-FLEX-tdTomato-T2A-SypEGFP-WPRE was a gift from Hongkui Zeng (Addgene viral prep # 51509-AAV1; http://n2t.net/addgene:51509; RRID:Addgene_51509). pENN.AAV.hSyn.Cre.WPRE.hGH was a gift from James M. Wilson (Addgene viral prep # 105553-AAV1; http://n2t.net/addgene:105553; RRID:Addgene_105553). pAAV-Ef1a-fDIO mCherry was a gift from Karl Deisseroth (Addgene viral prep # 114471-AAVrg; http://n2t.net/addgene:114471; RRID:Addgene_114471). pAAV-EF1a-Flpo was a gift from Karl Deisseroth (Addgene viral prep # 55637-AAV1; http://n2t.net/addgene:55637; RRID:Addgene_55637). pAAV-hSyn Con/Fon hChR2(H134R)-EYFP was a gift from Karl Deisseroth (Addgene viral prep # 55645-AAV8; http://n2t.net/addgene:55645; RRID:Addgene_55645). pAAV-EF1a-double floxed-hChR2(H134R)-EYFP-WPRE-HGHpA was a gift from Karl Deisseroth (Addgene viral prep # 20298-AAV5; http://n2t.net/addgene:20298; RRID:Addgene_20298). pAAV-EF1a-double floxed-hChR2(H134R)-mCherry-WPRE-HGHpA was a gift from Karl Deisseroth (Addgene viral prep # 20297-AAV5; http://n2t.net/addgene:20297; RRID:Addgene_20297). pAAV-EF1a-Cre was a gift from Karl Deisseroth (Addgene viral prep # 55636-AAVrg; http://n2t.net/addgene:55636; RRID:Addgene_55636). pAAV-nEF-Con/Fon-NpHR3.3-EYFP was a gift from Karl Deisseroth & INTRSECT 2.0 Project (Addgene viral prep # 137152-AAV8; http://n2t.net/addgene:137152; RRID:Addgene_137152). v237 was constructed by Dr. Hendrik Wildner, Institute of Pharmycology & Toxicology, University of Zurich, Switzerland. pAAV-hSyn-FLEX-TeLC-P2A-dTomato was a gift from Sandeep Datta (Addgene plasmid # 159102; http://n2t.net/addgene:159102; RRID:Addgene_159102)

